# Auto-STEED: A data mining tool for automated extraction of experimental parameters and risk of bias items from *in vivo* publications

**DOI:** 10.1101/2023.02.24.529867

**Authors:** Wolfgang Emanuel Zurrer, Amelia Elaine Cannon, Ewoud Ewing, Marianna Rosso, Daniel S. Reich, Benjamin V. Ineichen

**Author notes:** Equal contribution (Wolfgang Emanuel Zurrer, Amelia Elaine Cannon). **Correspondence to:** Benjamin Victor Ineichen, University of Zurich, Center for Reproducible Science, Zurich, Switzerland.

## Abstract

**Background:** Systematic reviews, i.e., research summaries that address focused questions in a structured and reproducible manner, are a cornerstone of evidence-based medicine and research. However, certain systematic review steps such as data extraction are labour-intensive which hampers their applicability, not least with the rapidly expanding body of biomedical literature.

**Objective:** To bridge this gap, we aimed at developing a data mining tool in the R programming environment to automate data extraction from neuroscience *in vivo* publications. The function was trained on a literature corpus (n=45 publications) of animal motor neuron disease studies and tested in two validation corpora (motor neuron diseases, n=31 publications; multiple sclerosis, n=244 publications).

**Results:** Our data mining tool Auto-STEED (Automated and STructured Extraction of Experimental Data) was able to extract key experimental parameters such as animal models and species as well as risk of bias items such as randomization or blinding from *in vivo* studies. Sensitivity and specificity were over 85 and 80%, respectively, for most items in both validation corpora. Accuracy and F-scores were above 90% and 0.9 for most items in the validation corpora. Time savings were above 99%.

**Conclusions:** Our developed text mining tool Auto-STEED is able to extract key experimental parameters and risk of bias items from the neuroscience *in vivo* literature. With this, the tool can be deployed to probe a field in a research improvement context or to replace one human reader during data extraction resulting in substantial time-savings and contribute towards automation of systematic reviews. The function is available on Github.

## 1. Introduction

Synthesising evidence is an essential part of scientific progress (1). To this end, systematic reviews— i.e. the rigorous identification, appraisal, and integration of all available evidence on a specific research question—have become a default tool in clinical research (2). Yet, they are also increasingly employed for preclinical *in vivo* research (3-6).

Systematic reviews allow the identification of trends that may be missed when reviewing individual, smaller studies, and add soundness to one’s conclusions. For this reason, the use of systematic reviews in animal research is an acknowledged aid to implementing the reduction, replacement, and refinement of animal experiments (7), e.g., by gaining knowledge without the use of new animal experiments or by improving the ethical position of animal research by increasing the value and reliability of research findings (8). Additionally, the practice of systematic reviews fosters a culture of transparent, reproducible, and rigorous scientific practice, pivotal and necessary in ensuring a responsible use of animals in research.

Despite the importance of systematic reviews, the process of manual evidence synthesis is highly laborious (9). This problem is further hampered by the skyrocketing amount of publications in the biomedical field: over 1 million papers pour into PubMed each year (10), and these numbers are set to increase still further in the near future (11). With this, it becomes increasingly difficult to keep abreast with the published evidence which in turn precludes evidence-based research (12). Thus, automation of systematic reviews is warranted to optimize the value of published data in the age of information overload. One particularly labour-intensive systematic review task which would profit from automation is data extraction (13, 14), i.e., the manual pulling of specific data from publications. Based on these shortcomings, we set out to develop a text mining tool to automatically extract key study parameters from publications of animal research modelling motor neuron diseases and multiple sclerosis. Our endeavour is focused on two key domains of experimental science, that is 1) disease model parameters such as animal models and species as well, and 2) risk of bias measures such as randomization or blinding.

## 2. Methods

### 2.1. Study protocol

The development of the text mining tool was part of a systematic review on neuroimaging findings in motor neuron disease animal models registered as prospective study protocol in the International Prospective Register of Systematic Reviews (PROSPERO, CRD42022373146, https://www.crd.york.ac.uk/PROSPERO/).

### 2.2. Literature corpora

Three literature corpora were included in this study: one for the training of the text mining toolbox and two for its validation. The training corpus was identified by searching Medline via PubMed for animal motor neuron disease models using the search string: *“motor neuron disease” OR motor neuron diseases [MeSH] OR “amyotrophic lateral sclerosis” OR “ALS” OR “MND” OR “SOD”* and limiting the search to the publication year 2021. The two validation corpora are derived from two in-house systematic reviews: a systematic review on neuroimaging findings in motor neuron disease animal models (PROSPERO-No: CRD42022373146, manuscript submitted) and a systematic review on neuroimaging findings in multiple sclerosis animal models (15) (PROSPERO-No: CRD42019134302).

### 2.3. Development of text mining tool

We defined items of interest to extract *a priori* which belong to two domains: first, experimental parameters including 1) animal sex, 2) animal species, 3) model disease, 4) number of experimental animals used, and 5-7) experimental outcomes, i.e., whether a respective study assessed behavioral, histological, or neuroimaging outcomes; second, risk of bias items including: 1) implementation in the experimental setup of any measure of randomization, 2) any measure of blinding, 3) prior sample size calculation (power calculation), 4) statement of whether conducted animal experiments are in accordance with local animal welfare guidelines, 4) statement of a potential conflict of interest, and 5) accordance with the ARRIVE guidelines (16). This second domain also includes an item for the data availability statement, i.e., a statement whether and where primary study data are available. Phrases associated with these parameters were systematically collected and integrated in a regular expression-based function using the R programming environment.

Performance of our text mining function was gauged using the following measures:

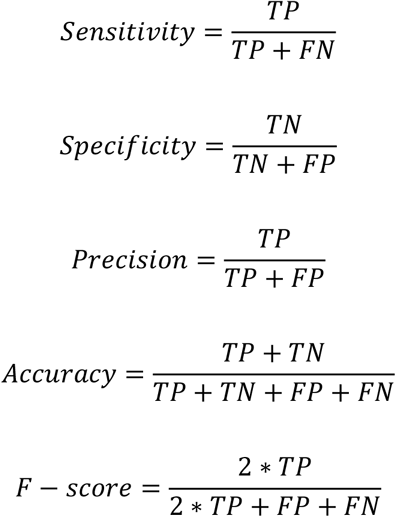

With TP, TN, FP, and FN being true positive, true negative, false positive, and false negative, respectively.

All included literature corpora have undergone dual and independent manual extraction of these parameters (WEZ, AEC, BVI) constituting the “gold standard” for data extraction. Mean extraction time was measured for both the human and the automated extraction to gauge time savings by the automated extraction. As defined in the protocol, for development of the text mining function in the training set, automated extraction of individual items was considered to be sufficiently accurate if they attained a sensitivity of 85% and a specificity of 80% (i.e., with a slightly higher sensitivity as per recommendation by the Systematic Living Information Machine [SLIM] consortium).

## 3. Results

### 3.1. General characteristics of literature corpora

We included three literature corpora with manual human annotation by two trained and independent reviewers. The training corpus comprised 45 individual publications on motor neuron disease animal models from 2021. The validation sets comprised 31 publications on neuroimaging in motor neuron disease animal models and 244 publications on neuroimaging in multiple sclerosis animal models with median publication years 2014 and 2009, respectively.

Median reporting prevalence of experimental parameters was 84%, 95%, and 95% in the training and in the two validation corpora, respectively. Median reporting prevalence of risk of bias items was 58%, 23%, and 25% in the training and in the two validation corpora, respectively. A detailed summary of literature corpora characteristics and reporting prevalence is presented in **Table 1**.

**Table 1:** Characteristics of included literature corpora and reporting prevalence for parameters to extract.

|  | Training corpus | Validation corpus 1 | Validation corpus 2 |
| --- | --- | --- | --- |
| Characteristics of eligible publications |  |  |  |
| Topic | Motor neuron disease animal models | Neuroimaging in motor neuron disease animal models | Neuroimaging in multiple sclerosis animal models |
| Number of publications | 45 | 31 | 244 |
| Publication year median and range | 2021 (2021-2021) | 2014 (2004 – 2020) | 2009 (1985 – 2017) |
| Number of different journals | 35 | 22 | 72 |
| Reporting prevalence |  |  |  |
| <u>Experimental parameters:</u> |  |  |  |
| Species | 100% | 100% | 100% |
| Sex | 87% | 61% | 88% |
| Model | 100% | 100% | >99% |
| Outcome histology | 80% | 90% | 85% |
| Outcome behaviour | 73% | 42% | 61% |
| Outcome imaging | 0% | 100% | 100% |
| <u>Risk of bias items:</u> |  |  |  |
| Randomization | 58% | 23% | 80% |
| Blinding | 53% | 29% | 33% |
| Animal welfare | 98% | 90% | 78% |
| Conflict of interest | 98% | 58% | 25% |
| Sample size calculation | 29% | 16% | <1% |
| ARRIVE guidelines | 29% | 0% | 1% |
| Data availability | 69% | 19% | 2% |

### 3.2. Architecture of text mining tool

Due to copyright restrictions for data mining from HTML, the tool was developed to extract data at PDF level of publications. First, the text mining function reads in and converts PDFs of respective publications to text. The text is then cleaned from certain keywords such as “random primer” reducing false positives for certain items to extract, e.g., randomization. Subsequently, the manuscript body is parsed into different sections (e.g., *abstract, introduction*, or *materials and methods*). This parsing is conducted based on the appearance of certain regular expressions (RegEx) such as “materials and methods”. Then, specific paper sections are mined for certain regular expressions based on RegEx libraries for each individual item to extract. The mining pipeline is depicted in **Figure 1**. The tool is available on Github: https://github.com/Ineichen-Group.

**Figure 1:**
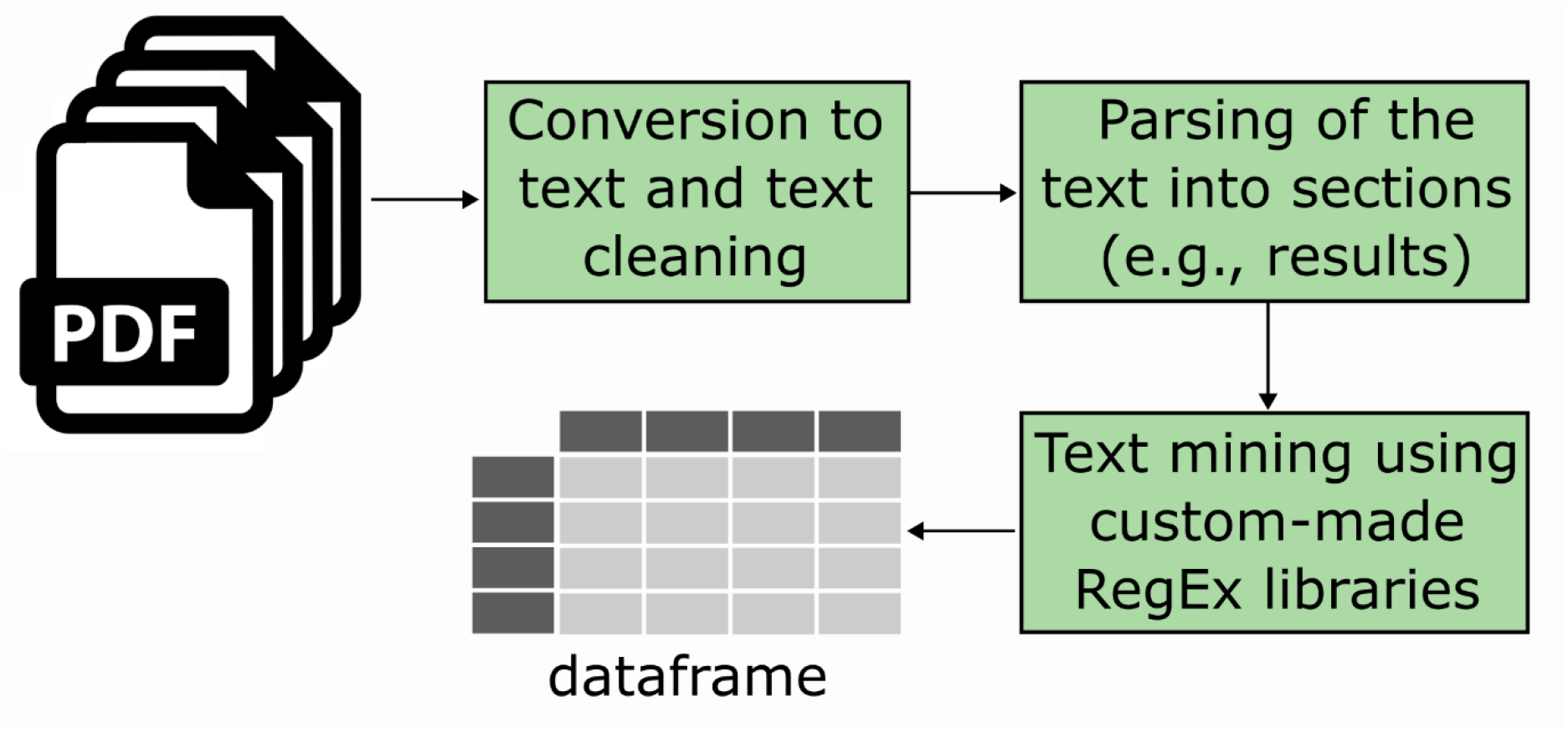
Architecture of the text mining function. PDFs of full texts are imported into the R environment, converted to text, and cleaned. Subsequently, the text is parsed into different sections such as “materials and methods” or “results”. Then, individual items to mine are extracted using custom-made Regex libraries and a data frame with the extracted items is created.

### 3.3. Accuracy

In the training set, the text mining function was tuned until a sensitivity of 85% and a specificity of 80% was reached for each individual item. The specificity threshold was not attained for the items “sample size calculation”, “sex”, and “outcome behaviour” with only 78%, 67% and 50%, respectively but with above-threshold sensitivity. Some items such as accordance with the ARRIVE guidelines or whether a conflict-of-interest statement was included reached a sensitivity close to 100%. F-scores and accuracy were above 90% for most items (**Table 2**).

**Table 2:** Summary of performance measures of RegEx compared with manual human ascertainment.

|  | Specificity | Sensitivity | Precision | Accuracy | F-score |
| --- | --- | --- | --- | --- | --- |
| **Training corpus (motor neuron diseases, n=45)** |  |  |  |  |  |
| Species | NA | 96 | 100 | 96 | 0.98 |
| Sex | 67 | 85 | 94 | 82 | 0.89 |
| Disease model | NA | 96 | 100 | 96 | 0.98 |
| Outcome histology | 89 | 92 | 97 | 91 | 0.94 |
| Outcome behaviour | 50 | 97 | 84 | 84 | 0.90 |
| Outcome imaging | 96 | NA | NA | 96 | NA |
| Randomization | 84 | 96 | 89 | 91 | 0.93 |
| Blinding | 95 | 92 | 96 | 93 | 0.94 |
| Animal welfare | NA | 86 | 97 | 84 | 0.92 |
| Conflict of interest | 100 | 98 | 100 | 97 | 0.99 |
| Sample size calculation | 0.78 | 92 | 63 | 82 | 0.75 |
| ARRIVE guidelines | 100 | 100 | 100 | 100 | 1.00 |
| Data availability | 85 | 94 | 94 | 91 | 0.94 |
| **Validation corpus 1 (motor neuron diseases, n=31)** |  |  |  |  |  |
| Species | NA | 100 | 100 | 100 | 1.00 |
| Sex | 100 | 74 | 100 | 84 | 0.85 |
| Disease model | NA | 90 | 100 | 90 | 0.95 |
| Outcome histology | 100 | 96 | 100 | 97 | 0.98 |
| Outcome behaviour | 78 | 85 | 76 | 81 | 0.79 |
| Outcome imaging | NA | 100 | 100 | 100 | 1.00 |
| Randomization | 100 | 86 | 100 | 97 | 0.92 |
| Blinding | 100 | 89 | 100 | 97 | 0.94 |
| Animal welfare | 100 | 89 | 100 | 90 | 0.94 |
| Conflict of interest | 92 | 94 | 94 | 94 | 0.94 |
| Sample size calculation | 81 | 80 | 44 | 81 | 0.57 |
| ARRIVE guidelines | 100 | NA | NA | 100 | NA |
| Data availability | 96 | 83 | 83 | 94 | 0.83 |
| **Validation corpus 2 (multiple sclerosis, n=244)** |  |  |  |  |  |
| Species | NA | 75 | 100 | 75 | 0.86 |
| Sex | 76 | 83 | 93 | 82 | 0.88 |
| Disease model | NA | 87 | 100 | 88 | 0.93 |
| Outcome histology | 64 | 96 | 93 | 91 | 0.95 |
| Outcome behaviour | 66 | 91 | 81 | 82 | 0.86 |
| Outcome imaging | NA | 94 | 100 | 94 | 0.97 |
| Randomization | 93 | 81 | 75 | 90 | 0.78 |
| Blinding | 98 | 85 | 96 | 93 | 0.90 |
| Animal welfare | 86 | 80 | 95 | 82 | 0.87 |
| Conflict of interest | 96 | 97 | 90 | 97 | 0.93 |
| Sample size calculation | 94 | 100 | 27 | 97 | 0.43 |
| ARRIVE guidelines | 100 | 100 | 100 | 100 | 1.00 |
| Data availability | 100 | 80 | 80 | 100 | 0.80 |

Specificity, sensitivity, precision, and accuracy are denoted in percentage. For details regarding
measures, please see the materials and methods section.

The mining function performed well on both validation corpora. In the motor neuron disease corpus, the mining function accomplished above-threshold specificity and sensitivity for most items, except for “outcome behaviour” with slightly below-threshold specificity and “data availability”, “sample size calculation”, and “sex” with slightly below-threshold sensitivity. In the multiple sclerosis validation corpus, additional items did not reach the specificity and sensitivity thresholds. However, F-scores and accuracy were above 90% for most items in the motor neuron disease validation corpus and above 80% in the multiple sclerosis corpus, respectively (**Table 2**).

### 3.4. Time savings automated versus manual extraction

Mean time for the manual extraction was 12 (± 8), 13 (± 7), and 15 (± 11) minutes per publication and per human reader for the training corpus and the two validation corpora, respectively. This amounts to a total of 540, 403, and 3660 minutes for one reader for the three corpora, respectively. In contrast, the mining function required 0.3 seconds to mine one record amounting to 0.23, 0.15, and 1.22 minutes for the three corpora. With this, the text mining function provides time savings above 99%.

### 3.5. Reporting of items on abstract versus full text level

For the experimental parameters, we quantified how commonly the respective items were reported in the abstract in addition to the full text. Disease models and species as well as outcome measures were commonly reported on abstract level in all three literature corpora with reporting frequencies between 95 – 100%. However, animal sexes were only rarely reported with reporting frequencies between 0 and 5%.

## Discussion

### Main findings

We developed *Auto-STEED* (Automated and STructured Extraction of Experimental Data), a text mining tool able to automatically extract key experimental parameters such as animal models and species as well as risk of bias items such as randomization or blinding from preclinical *in vivo* studies. The function shows a high sensitivity, specificity, and accuracy for most items to extract in two validation literature corpora, one in a similar field like the training corpus (motor neuron diseases) and one in a different field (multiple sclerosis) and both including older publications. Using this approach, time savings to extract these items are above 99%. We also show that mining from abstracts instead of full texts would be feasible for certain key experimental parameters.

### Findings in the context of existing evidence

Our developed text mining tool performs well on literature corpora outside of the field they have been developed in as well as in corpora with older median publication years. The tool has been developed in a literature corpus dealing with motor neuron disease animal models and only comprising publications from 2021. In contrast, one of the validation literature corpora was in the field of multiple sclerosis animal models and had a median publication year 2009 (with some papers going back to 1985). And although the accuracy was slightly lower in this literature corpus, this shows that reporting of experimental parameters and risk of bias items is similar between neuroscience subfields. Thus, our developed function could be applied to literature bodies of other research fields.

Despite its high accuracy, our model is not yet at a level appropriate for the evaluation of individual publications. Thus, it will not fully replace human extraction. However, such an automated approach has two potential fields of application: first, it is considered suitable for deployment on larger reference libraries (>1000 records) in a research-improvement context (17) and/or to probe a certain field or literature bodies for risk of bias and key experimental parameters. Second, such a method could be deployed to replace one human reader which would still save a substantial amount of labour (14, 18). Human-machine disagreements could be checked manually.

Similar approaches have been leveraged to extract specific information—such as the study population, intervention, outcome measured and risks of bias—from abstracts (19) or full texts (17, 20). Bahor and colleagues developed a text mining function in a literature body of stroke animal models able to extract certain risk of bias items including randomization, blinding, and sample size calculation (21). The achieved accuracy was between 67-86% for randomization (our approach: 90-97%), 91-94% for blinding (our approach: 93-97%), and 96-100% for sample size calculation (our approach: 81-97%). With this, our developed tool has a similar accuracy scope and does complement former tool by extracting additional risk of bias items such as statement of a conflict of interest, accordance with local animal welfare regulations, a data availability statement, and accordance with the ARRIVE guidelines (16). Another text mining toolbox underpinned by natural language processing (NLP) was developed by Zeiss and colleagues (19): This toolbox extracts data such as species, model, genes, or outcomes from PubMed abstracts with F-scores between 0.75 and 0.95.

For many tasks, NLP models seem to consistently outperform RegEx-based text mining (22). Yet they are more complex and labour-intensive to develop and thus only warrant application in more complex extraction tasks. Wang and colleagues tested performance of a variety of models such as convolutional neural networks to extract risk of bias items from preclinical studies (17). These models significantly outperformed RegEx-based methods for four risk of bias items with F-scores between 0.47-0.91. The validity of NLP for such tasks has also been corroborated by SciScore—a proprietary NLP tool that can automatically evaluate the compliance of publications with six rigour items taken from the MDAR framework and other guidelines (20). These items mostly relate to risk of bias, including compliance with animal welfare regulations, blinding/randomisation, prior sample size calculation and other items such as organism or sex. SciScore was developed on a training corpus from PubMed open access articles. In contrast, our approach was developed on preclinical neuroscience corpora thus being more tailored to this field.

Although we initially aimed to also extract used animal numbers from publications, we had to abandon this goal due to a highly unstandardized nature of reporting, i.e., in methods/results section, in tables, in figure legends, in graphs or only separately reported for different experimental and control groups. One potential solution to this problem could be to consider this as an NLP categorisation task with small (e.g., n<10 animals), medium (n=10-50 animals) and large (n>100 animals) studies.

### Limitations

First, our approach has been developed and tested in the realm of preclinical neuroscience. It is currently not clear how well the tool would perform in fields outside of neuroscience research, e.g., in the preclinical cancer literature. Second, our approach requires full-text PDFs for mining. Mining in online publication versions, i.e., on HTLM would mitigate certain issues associated with converting a PDF into text including unstandardized PDF layouts and paper sections per journal. However, although text mining will be exempted from copyright restrictions in the EU within the coming years (23), expensive licences are still required to mine online versions of publications.

## Conclusions

Our developed text mining tool Auto-STEED is able to extract key risk of bias items and experimental parameters from the neuroscience *in vivo* literature. Accelerating the usually labour-intensive data extraction during a systematic review is an important contribution towards automation of systematic reviews.

## Glossary

NLP: natural language processing
RegEX: regular expressions

## Acknowledgments

We thank Robert Wyatt from Matching Mole for help with data analysis.

## Data availability

The text mining function is freely available on Github: https://github.com/Ineichen-Group

## Author contributions

Conception and design of study: EE, BVI

acquisition of data: WEZ, AEC, EE, BVI

analysis of data: WEZ, AEC, BVI

drafting the initial manuscript: BVI

all authors critically revised the paper draft.

